## supplementary information for "Demographic inference for spatially heterogeneous populations using long shared haplotypes"

June 9, 2023

### S.1 Random walk approximation of the skew diffusion

Fix $m_+ \in (0,1)$, $m_- \in (0,1)$ and $L \in 2\mathbb{N}+1$. Consider a random walk $(Y_n^{(L)}, n \in \mathbb{N})$ taking values in $\mathcal{X}_L := [\![ -\frac{L+1}{2}, \frac{L+1}{2} ]\!]^2$, with transition kernel given by

$$\Pi_{x,y} := \begin{cases} \frac{m_\alpha}{4} & \text{for } |x-y| = 1 \text{ and either } \alpha x^{(1)} > 0 \text{ or } \alpha y^{(1)} > 0, \\ \frac{m_+ + m_-}{8} & \text{for } |x-y| = 1 \text{ and } x^{(1)} = y^{(1)} = 0, \\ 1 - \sum_{z:|z-x|=1} \Pi_{x,z} & \text{for } y = x. \end{cases}$$

This kernel is depicted on Figure S.1. The following is then a direct consequence of [IP16, Theorem 1.1].

**Theorem 1.** *Suppose that $(L_n, n \in \mathbb{N})$ and $(\delta_n, n \in \mathbb{N})$ are two sequences such that $\delta_n \to 0$ and $\delta_n L_n \to \infty$ as $n \to \infty$, and fix $r > 0$. Set*

$$Y_t^n := \delta_n Y^{(L_n)}_{rt/\delta_n^2}, \quad t \geq 0.$$

*Then, as $n \to \infty$, $(Y_t^n, t \geq 0)$ converges in distribution to $(X_t, t \geq 0)$, solution of (6), with $\sigma_\alpha^2 = r\frac{m_\alpha}{2}$.*

Given a set of sampling positions, we thus chose a step size $\delta$ and a grid size $L$ such that $\delta$ is small compared to the mean distance between samples and $L$ is large compared to the maximum distance between the samples. We then choose $m_+$, $m_-$ and $r$ as functions of $\sigma_+$ and $\sigma_-$, and we approximate $P_t\phi(x)$ by

$$\mathbb{E}_x \left[ \phi(\delta\, Y^{(L)}_{t/\delta^2}) \right]$$

in (11). Note that if $\sigma_\alpha^2 < 1/2$ for $\alpha \in \{+,-\}$, then we can take $r = 1$, $m_\alpha = 2\sigma_\alpha^2$ above, but if $\sigma_\alpha^2 \geq 1/2$ for some $\alpha$, then we choose $r > 0$ large enough that $\sigma_\alpha^2/r < 1/2$ for $\alpha \in \{+,-\}$ and we take $m_\alpha = 2\sigma_\alpha^2/r$.

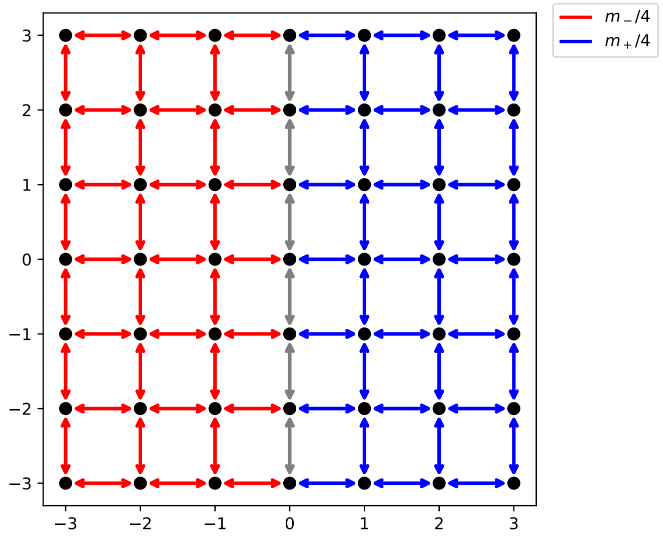


Figure S.1: Transition kernel of the random walk $(Y_t^{(L)}, t \geq 0)$ used to approximate the transition density of the skew diffusion. At each step, the probability of jumping along each edge depends on the color of the edge ($m_-/4$ for red edges, $m_+/4$ for blue edges, and $(m_+ + m_-)/8$ for grey edges).

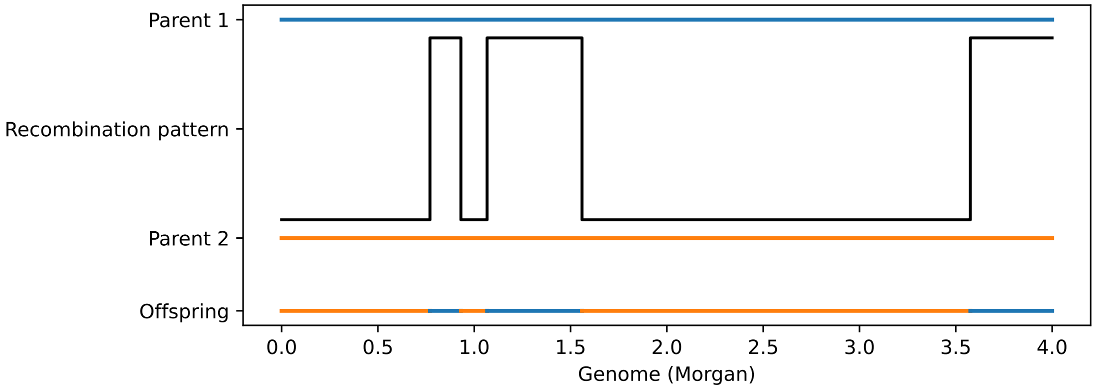


Figure S.2: Example of a recombination pattern and the resulting offspring genome. The crossover take place at the points of a standard Poisson process along the genome, and the parent genome that is ancestral to that of the new genome changes at each of these points.

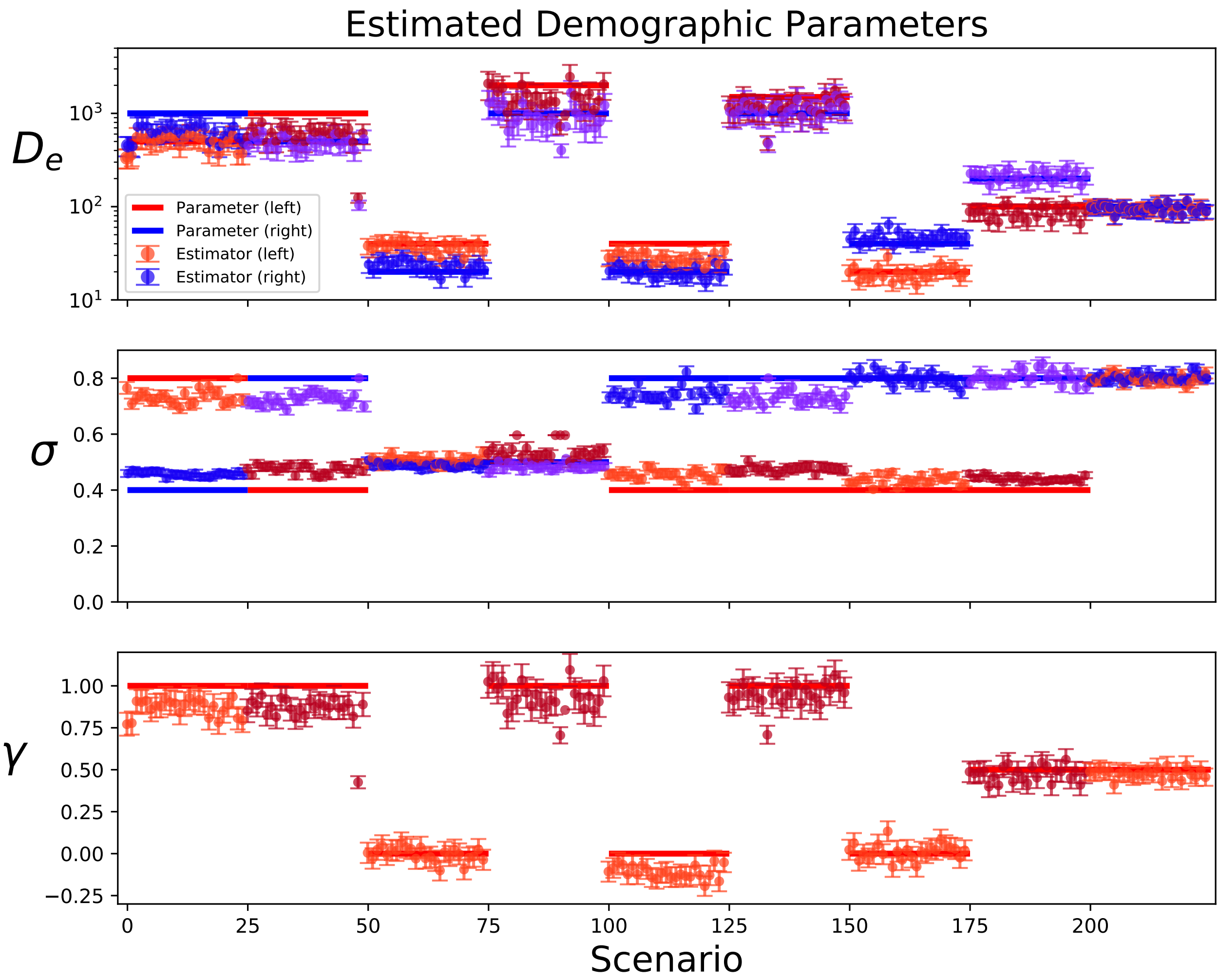


Figure S.3: Simulated (straight lines) vs. estimated (dots) parameters for 20 independent runs for each of 9 scenarios. Diffusion and effective neighbourhood size on the left of the interface are shown in red, and those on the right are shown in blue. The error bars correspond to the 95% confidence interval obtained from the Fisher information matrix (see the discussion in the main text). The parameter $\gamma$ is the growth exponent of the population density (it is the same on both sides of the interface), such that the effective neighbourhood size $t$ generations in the past is $D_e t^{-\gamma}$.

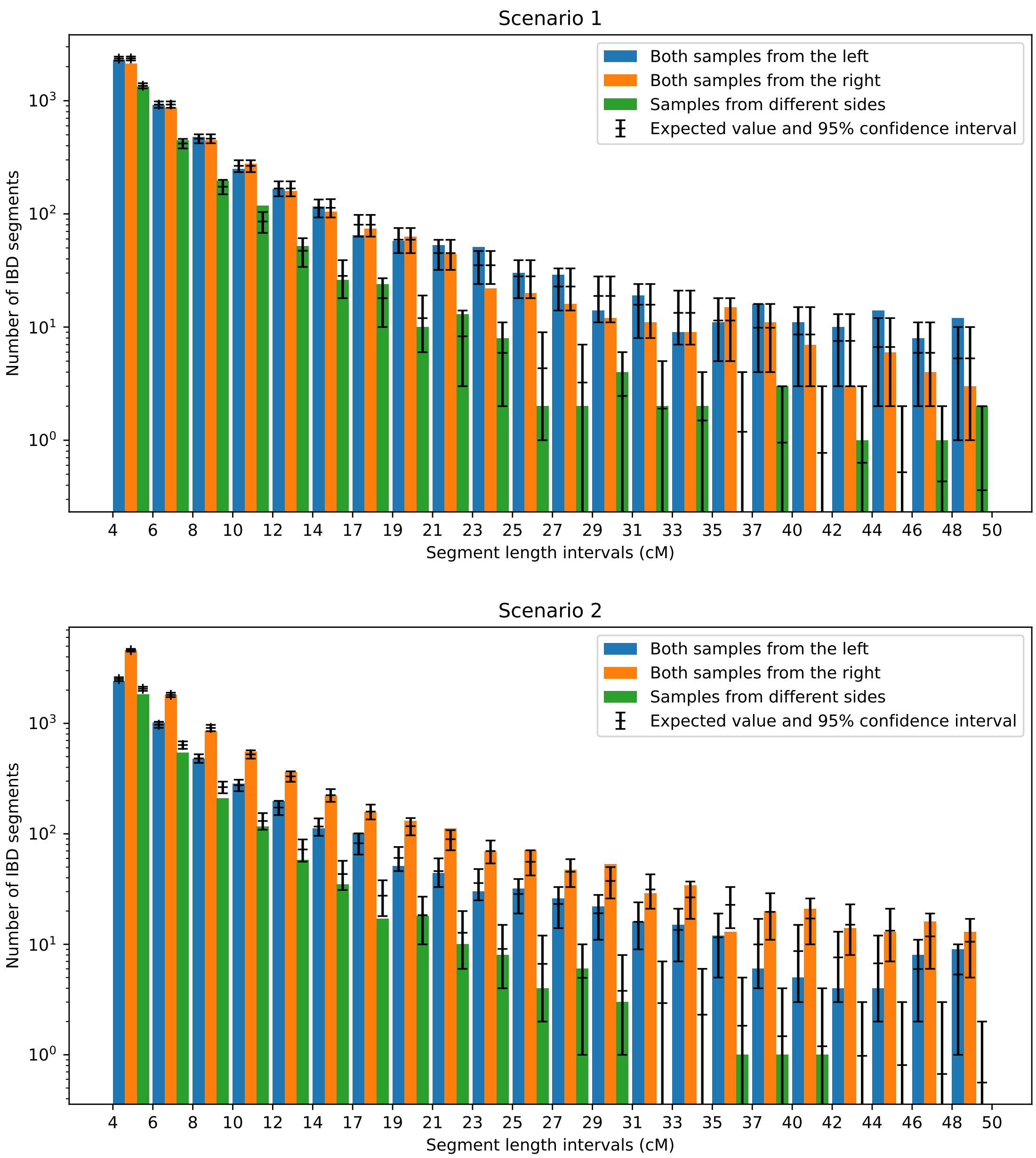


Figure S.4: Number of IBD segments of different lengths observed in one simulation of the SLFV for scenarios 1 and 2 (see Table 1). The bar plot shows the sum of all observed IBD segments of each length interval for all pairs of individuals respectively both on the left side of the interface, both on the right side, and on different sides. The expected value computed with (11) with the simulation parameters and the associated 95% confidence interval of the Poisson distribution are shown in black for each length interval and each sampling configuration.

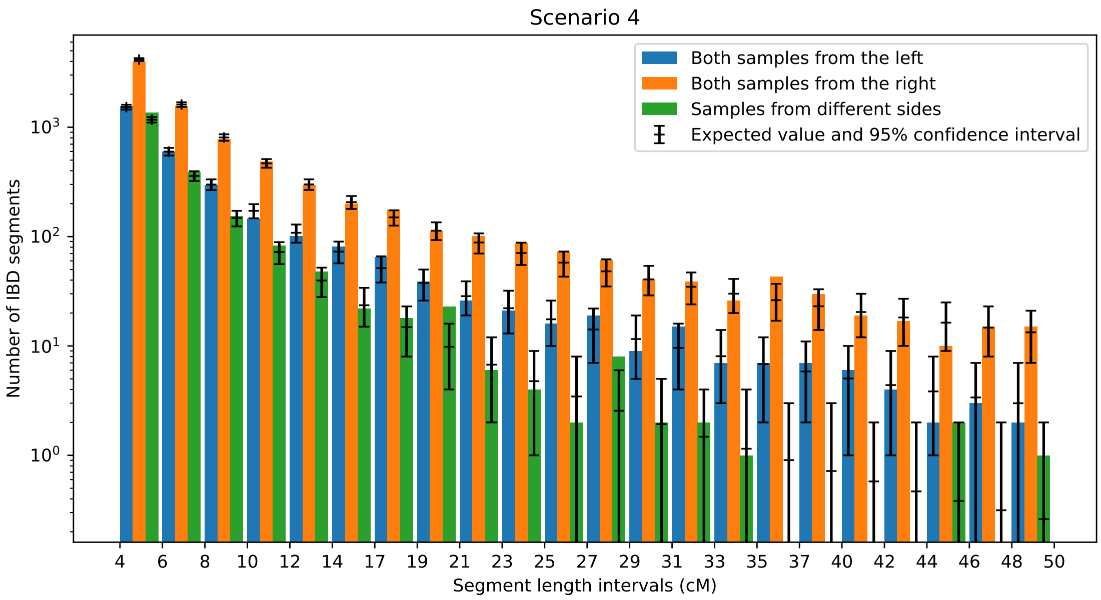


Figure S.4: Corresponding graphic for scenario 4 (see Table 1).

### S.2 Proof of Lemma B.7

*Proof of Lemma B.7.* We start by proving that $(Y(t), t \geq 0)$ is ergodic with respect to the uniform probability measure on $[-R, R]$. Let $\mathcal{L}^Y$ denote the infinitesimal generator of $(Y(t), t \geq 0)$. For $f$ and $g$ two bounded and measurable functions on a set $B \subset \mathbb{R}$, let

$$\langle f, g \rangle_B := \int_B f(x) g(x) dx.$$

We want to show

$$\langle \mathcal{L}^Y f, g \rangle_{[-R,R]} = \langle f, \mathcal{L}^Y g \rangle_{[-R,R]}. \tag{S.1}$$

Recall the definition of the operator $E$ in (B.8) and note that, $\mathcal{L}^Y f = \mathcal{L} E f$ on $[-R, R]$, where $\mathcal{L}$ is defined in (3). In addition, since $\Phi(x, y) = \Phi(y, x)$, for any $f, g \in L^2(\mathbb{R})$,

$$\langle \mathcal{L} f, g \rangle_{\mathbb{R}} = \langle f, \mathcal{L} g \rangle_{\mathbb{R}}.$$

However, $Ef \notin L^2(\mathbb{R})$. To circumvent this, for $A \geq R$, define

$$\Phi^A(x, y) := \begin{cases} \Phi(x, y) & \text{if } |x| \leq A \text{ and } |y| \leq A, \\ 0 & \text{otherwise.} \end{cases}$$

Further let $(\xi^A_t, t \geq 0)$ be a random walk on $\mathbb{R}$ with generator

$$\mathcal{L}^A f(x) := \int_{\mathbb{R}} \Phi^A(x, y)(f(y) - f(x)) dy \tag{S.2}$$

which coincides with $(\xi_t, t \geq 0)$ up to the random time $T_A := \inf\{t \geq 0 : |\xi_t| > A\}$. Finally, for $|x| \leq A$, define

$$E^A f(x) := \mathbb{E}_x \left[ f(\xi^A_{T^A_0}) \right], \quad \text{with} \quad T^A_0 := \inf\{t \geq 0 : \left|\xi^A_t\right| \leq r_+\}. \tag{S.3}$$

Then the operator $\mathcal{L}^A$ is self-adjoint in $L^2([-A, A])$ and

$$\mathcal{L}^A E^A f(x) = 0 \qquad \text{for } R < |x| \leq A, \tag{S.4}$$

$$E^A f(x) = f(x) \qquad \text{for } |x| \leq R. \tag{S.5}$$

As a result, for $f, g : [-R, R] \to \mathbb{R}$ bounded and measurable,

$$\langle \mathcal{L}^A E^A f, E^A g \rangle_{[-A,A]} = \langle E^A f, \mathcal{L}^A E^A g \rangle_{[-A,A]}.$$

By (S.4) and (S.5), this is

$$\langle \mathcal{L}^A E^A f, g \rangle_{[-R,R]} = \langle f, \mathcal{L}^A E^A g \rangle_{[-R,R]}.$$

It thus remains to let $A \to \infty$. First note that, for $A$ large enough, $\mathcal{L}^A f(x) = \mathcal{L} f(x)$ for all $x \in [-R, R]$. Furthermore, since $T_A \to \infty$ as $A \to \infty$ almost surely, $\xi^A_{T^A_0} \to \xi_{\alpha(0)}$ as $A \to \infty$ almost surely. Hence, by dominated convergence, for all $x \in \mathbb{R}$ and for all bounded and measurable $f$,

$$E^A f(x) \underset{A \to \infty}{\longrightarrow} E f(x). \tag{S.6}$$

We thus obtain

$$\langle \mathcal{L} E f, g \rangle_{[-R,R]} = \langle f, \mathcal{L} E g \rangle_{[-R,R]},$$

which is (S.1). As a result the uniform measure on $[-R, R]$ is invariant for $Y$. The fact that $(Y(t), t \geq 0)$ is ergodic then follows from the form of its generator, noting that $\mathcal{L}^Y f \equiv 0$ implies that $f$ is almost everywhere constant. □

Before proving the second part of Lemma B.7, let us show the following.

**Lemma S.2.1.** *For any $f : [-R, R] \to \mathbb{R}$ bounded and measurable,*

$$\int_{-\infty}^{R} \int_{R}^{+\infty} \Phi(x, y) \left(Ef(x) - Ef(y)\right) dydx = 0$$

*and*

$$\int_{-R}^{+\infty} \int_{-\infty}^{-R} \Phi(x, y) \left(Ef(x) - Ef(y)\right) dydx = 0$$

*Proof.* Recall the definition of $\mathcal{L}^A$ and $E^A$ in (S.2) and (S.3). By (S.4), for any $f : [-R, R] \to \mathbb{R}$ bounded and measurable,

$$\int_{R}^{A} \mathcal{L}^A E^A f(x) dx = 0.$$

Since $\mathcal{L}^A$ is self-adjoint on $[-A, A]$,

$$0 = \int_{-A}^{A} \int_{-A}^{A} E^A f(x) \Phi^A(x, y) \left(\mathbb{1}_{\{y>R\}} - \mathbb{1}_{\{x>R\}}\right) dydx.$$

But

$$\mathbb{1}_{\{y>R\}} - \mathbb{1}_{\{x>R\}} = \mathbb{1}_{\{y>R, x\le R\}} - \mathbb{1}_{\{y\le R, x>R\}}.$$

The above term is thus

$$\int_{-A}^{R} \int_{R}^{A} \Phi^A(x, y) E^A f(x) dydx - \int_{R}^{A} \int_{-A}^{R} \Phi^A(x, y) E^A f(x) dydx.$$

Since $\Phi^A(x, y) = \Phi^A(y, x)$, we obtain

$$\int_{-A}^{R} \int_{R}^{A} \Phi^A(x, y) \left(E^A f(x) - E^A f(y)\right) dydx = 0.$$

Letting $A \to \infty$ and using (S.6), we obtain the first statement of Lemma S.2.1. The second statement follows by a similar argument. □

We now finish the proof of Lemma B.7.

*Conclusion of proof of Lemma B.7.* Let us start by computing $\frac{1}{2R} \int_{-R}^{R} h^+(x) dx$. By the first part of Lemma B.7 and (B.7),

$$\frac{1}{2R} \int_{-R}^{R} h^+(x) dx = \frac{u}{2R} \int_{-R}^{R} \int_{-\infty}^{R} \Phi(x, y) \left(E\iota(y) - x\right) dydx + \frac{u}{2R} \int_{-R}^{R} \int_{R}^{+\infty} \Phi(x, y)(y - x) dydx.$$

Since $E\iota(y) = y$ when $|y| \le R$ and $\Phi(x, y) = 0$ when $|x - y| > 2R$, this is

$$\frac{1}{2R} \int_{-R}^{R} h^+(x) dx = \frac{u}{2R} \int_{-R}^{R} \int_{-R}^{R} \Phi(x, y) (y - x) dydx + \frac{u}{2R} \int_{-R}^{\infty} \int_{-\infty}^{-R} \Phi(x, y) \left(E\iota(y) - E\iota(x)\right) dydx$$
$$+ \frac{u}{2R} \int_{-\infty}^{R} \int_{R}^{+\infty} \Phi(x, y)(y - x) dydx.$$

The first term on the right hand side is zero because $\Phi(x,y) = \Phi(y,x)$ and the second term is zero by Lemma S.2.1. Replacing $y$ by $x+z$ in the last term, we have

$$\frac{1}{2R}\int_{-R}^{R} h^{+}(x)dx = \frac{u}{2R}\int_{-\infty}^{R}\int_{0}^{+\infty} \Phi(x,x+z)\mathbb{1}_{\{x+z>R\}} z dz dx.$$

But for $x+z>R$, $B(x+z,r_{-})\cap\mathbb{H}^{-}=\emptyset$ and

$$\Phi(x,x+z) = \frac{|B(x,R)\cap B(x+z,R)|}{V_R^2} = \Phi(R,R+z).$$

Furthermore, the expression above is zero when $z \geq 2R$. Changing the order of integration, we obtain

$$\begin{aligned}\frac{1}{2R}\int_{-R}^{R} h^{+}(x)dx &= \frac{u}{2R}\int_{0}^{2R} \Phi(R,R+z)z\int_{R-z}^{R} dxdz \\ &= \frac{u}{2R}\int_{0}^{2R} \Phi(R,R+z)z^2 dz \\ &= \frac{\sigma_{+}^{2}}{4R}.\end{aligned}$$

By the same argument, one arrives at

$$\frac{1}{2R}\int_{-R}^{R} h^{-}(x)dx = \frac{\sigma_{-}^{2}}{4R}.$$

□
